# Biogeographical history shapes evolution of reproduction in a global warming scenario

**DOI:** 10.1101/2022.02.25.481927

**Authors:** Marta A. Santos, Marta A. Antunes, Afonso Grandela, Ana Carromeu-Santos, Ana S. Quina, Mauro Santos, Margarida Matos, Pedro Simões

**Affiliations:** cE3c – Centre for Ecology, Evolution and Environmental Changes & CHANGE – Global Change and Sustainability Institute, Lisboa, Portugal; Departamento de Biologia Animal, Faculdade de Ciências, Universidade de Lisboa, Lisboa, Portugal; CESAM – Centre for Environmental and Marine Studies, Universidade de Aveiro and Faculdade de Ciências, Universidade de Lisboa, Lisboa, Portugal; Departament de Genètica i de Microbiologia, Grup de Genòmica, Bioinformàtica i Biologia Evolutiva (GBBE), Universitat Autònoma de Barcelona, Spain

**Keywords:** Drosophila, experimental evolution, global warming, historical background, thermal adaptation, thermal reaction norms

## Abstract

Adaptive evolution might be critical for animal populations to thrive on the fast-changing natural environments. Ectotherms are particularly vulnerable to global warming and, although their limited coping ability has been suggested, few real-time evolution experiments have directly accessed their evolutionary potential. Here, we report a long-term experimental evolution study addressing the evolution of thermal reaction norms, after ∼30 generations under different thermal environments. We analyzed the evolutionary dynamics of *Drosophila subobscura* populations as a function of the thermally variable environments in which they evolved and their distinct biogeographical background. Our results showed clear differences between the historically differentiated populations: while the northern *D. subobscura* populations showed a temporal increase in performance at higher temperatures, their southern counterparts presented the opposite pattern. This suggests that the northern populations might be better equipped to cope with the current rising temperatures. Remarkably, no effect of thermal selection was found. The lack of a clear long-term adaptive response at higher temperatures after evolution under a global warming scenario raises concerns about the evolutionary potential of ectotherms. Our results highlight the complex nature of thermal responses in face of environmental heterogeneity and emphasize the importance of considering intra-specific variation in thermal evolution studies.

## Introduction

Biodiversity is under great pressure due to rapid environmental changes associated with global warming. Adaptive evolution and plasticity may be important means for organisms to cope with the increasingly warmer temperatures (Chevin et al. 2010; Merilä et al. 2016). Plasticity can also be adaptive if, for instance, environmental cues occurring during development allow for animals to better cope with environmental conditions as adults (see reviews in (Kelly 2019; Rodrigues and Beldade 2020; Buckley and Kingsolver 2021). Ectotherms are particularly vulnerable animals as their body temperature is highly constrained by the environment, which has a major impact on their biological processes and, thus, on their fitness (Angilletta 2009; Hoffmann and Sgrò 2018). To make matters worse, recent evidence showed a limited ability of ectotherms to evolutionarily respond to upper temperature shifts (Kellermann and van Heerwaarden 2019; MacLean et al. 2019), as well as limited scope for plasticity in that environmental context (Gunderson and Stillman 2015; Sørensen et al. 2016).

A thermal adaptive response will, expectedly, lead to an optimal performance within the range of temperatures to which the organisms have most exposure, and lower fitness at more extreme values (Angilletta 2009). In the context of global warming, one might expect a shift in the optimum performance towards higher temperatures, provided genetic variation is present and other evolutionary constraints do not play a major role (Kellermann and van Heerwaarden 2019; Kristensen et al. 2020). These shifts can be observed through changes in the thermal reaction norms, which describe the population’s performance as a function of temperature range. According to Angilletta *et al*. (2003), different types of trade-offs may be involved in changes of elevation or shape of thermal reaction norms. On the one hand, the allocation of resources (*e*.*g*., energy differently allocated to two traits) or acquisition trade-offs (*e*.*g*., between maximizing the acquisition of resources *vs*. minimizing the risk of mortality) will mainly lead to shifts in trait means, *i*.*e*., to the elevation of the thermal reaction norm. On the other hand, the evolution under a specialist-generalist trade-off will expectedly lead to a change in shape, if an increase in performance at a given temperature causes a decrease at other temperatures. These different sources of variation of thermal reaction norms are not mutually exclusive. The genetic variation available for these different dimensions will govern the evolution of reaction norms (Huey and Kingsolver 1989; Angilletta et al. 2003; Berger et al. 2013). Generalist *vs*. specialist trade-offs can underlie patterns of local adaptation, even though adaptation costs are not always present (Hereford 2009). There is evidence of thermal plasticity suggestive of local adaptation in natural populations of ectotherms, with populations geographically distinct presenting different thermal reaction norms (Trotta et al. 2006; Berger et al. 2013; Austin and Moehring 2019; Klepsatel et al. 2019), but it is not universal (e.g., (Klepsatel et al. 2013; Clemson et al. 2016). It is, thus, critical to address how plasticity evolves in face of rapid thermal changes and whether this variation is consistent across populations, to better understand the consequences of the current global warming scenario.

Experimental evolution, the real-time study of populations across several generations, in highly controlled and reproducible conditions (Kawecki et al. 2012), is a very powerful approach to address the adaptation to novel environmental conditions, including the evolution of plasticity. It has been frequently used to characterize the tempo and mode of thermal evolution of ectotherms, namely in ecological scenarios associated with global warming (Schou et al. 2014; Manenti et al. 2015; Kellermann and van Heerwaarden 2019; Kinzner et al. 2019; Liukkonen et al. 2021). The native Palearctic *Drosophila subobscura* is an emblematic case study of ectotherm thermal adaptation. It shows local adaptation to distinct natural habitats and latitudinal clinal variation for chromosomal inversion frequencies in Europe, North America, and South America (Prevosti et al. 1988; Rezende et al. 2010). These karyotypic polymorphisms have been shown to change worldwide as a result of global warming (Balanyá et al. 2006) and to respond to sudden heat waves (Rodríguez-Trelles et al. 2013). Local adaptation in the reproductive performance of *D. subobscura* during heat stress has also been reported (Porcelli et al. 2017). This species presents clear plastic responses to thermal variation (Fragata et al. 2016), with a key role of developmental temperature in shaping adult reproductive performance (Simões et al. 2020; Santos et al. 2021a). The developmental temperature range of *D. subobscura* is 6 – 26ºC (Moreteau et al. 1997; David et al. 2005; Schou et al. 2017) with optimal viability and preferred body temperature between 16ºC and 20ºC (Rego et al. 2010; Castañeda et al. 2013; Schou et al. 2017). Shifts in this species’ thermal reaction norms have been observed for locomotor behavior (Mesas et al. 2021) and thermal tolerance (Schou et al. 2016; MacLean et al. 2019), although in the latter case mostly for lower thermal limits.

For the last three years, we have been performing a thermal experimental evolution study on historically differentiated *D. subobscura* populations, derived from the extreme latitudes of the European cline (Simões et al. 2017). These populations have been subjected to thermal selective regimes that differ in mean temperature and/or thermal amplitude (Santos et al. 2021b). We have previously found no evidence for evolutionary changes in thermal reaction norms after nine generations in circadian fluctuating or global warming-like environments (Santos et al. 2021b). This finding may result from a small number of elapsed generations or a general lack of genetic variation for response to thermal selection.

Here, we present an extended report of the evolutionary response of these populations, spanning ∼30 generations of thermal evolution, and thus testing a longer-term adaptive shift in thermal reaction norms. As in Santos et al. (2021), the thermal reaction norms were estimated through the population’s reproductive performance at three different temperatures (14ºC, 18ºC, and 24ºC). With this design, we aim to respond to three main questions: *(i)* Does longer-term evolution under thermally diverse environments change the populations’ thermal reaction norm? *(ii)* Do thermal reaction norms change between short- and longer-term evolution? *(iii)* Can we find the signature of biogeographical history in the populations’ thermal evolutionary response?

First and foremost, we expect that differential plasticity between thermal selection regimes may arise in a longer-term study, provided there is a correlated change in reaction norms due to adaptation to the dynamic thermal environments. Because of almost 30 generations experiencing a wide range of temperatures, we foresee a higher performance of the global warming-evolved populations in more extremal thermal environments. Second, we expect that fluctuating and warming-evolved populations to perform worse than the control lines in intermediate thermal conditions, assuming a trade-off between maximal performance and performance breath of the reaction norms (Huey and Kingsolver 1989; Angilletta et al. 2003). Finally, we think that geographical variation for plasticity may become apparent, driven by an interaction of thermal selection with the distinct genetic backgrounds.

## Material and Methods

### Population maintenance and thermal selection regimes

The experimental populations derived from collections from natural populations of *Drosophila subobscura* in two contrasting latitudes of the species’ European cline: Adraga, Portugal (38º48′ N) and Groningen, The Netherlands (53º13′ N). The laboratory populations (PT, from Portugal, and NL, from The Netherlands) were established in early September 2013 and three-fold replicated shortly after. All PT_1-3_ and NL_1-3_ populations were maintained in discrete generations with synchronous 28-day cycle, 12L:12D photoperiod, and constant 18ºC temperature (control conditions, C). The flies were reared in ∼30 mm^3^ glass vials with controlled densities in both eggs (70 eggs per vial) and adults (50 adults per vial). Egg collection for the following generation was around peak fecundity (seven to ten days old flies) and the census sizes per generation ranged between 500 and 1000 individuals (see (Simões et al. 2017), for more details).

Two new thermal selection regimes were derived in January 2019, by the time PT and NL populations had undergone 70 generations of lab evolution. The *circadian fluctuation* regime (F populations, FPT_1-3_ and FNL_1-3_) and the *global warming* regime (W populations, WPT_1-3_ and WNL_1-3_) – see (Santos et al. 2021b). The F populations are under a daily temperature regime that fluctuates between 15ºC and 21ºC, with a mean daily temperature of 18ºC, a cycle that is constant across generations. The W populations experience a daily fluctuation initially similar to the F lines but with a per generation increase of 0.18ºC in daily mean and 0.54ºC in daily amplitude (see the thermal profiles on Figure 1). This simultaneous increase in thermal mean and amplitude per generation was, to our knowledge, not yet tested in other experimental studies. Taking into account the global warming pace of 0.1-0.3ºC per decade (IPCC 2018), a mean increase of 0.18ºC per generation is equivalent to the thermal changes experienced in nature by an organism with a ∼10-year generation time. The progressively divergent upper and lower temperature extremes (increases of 0.44ºC in upper and 0.08ºC in lower limits, per generation) fit well with the projected 2-fold increase in thermal extremes relative to mean temperature in mid-latitude location (IPCC 2018).

**Figure 1.**
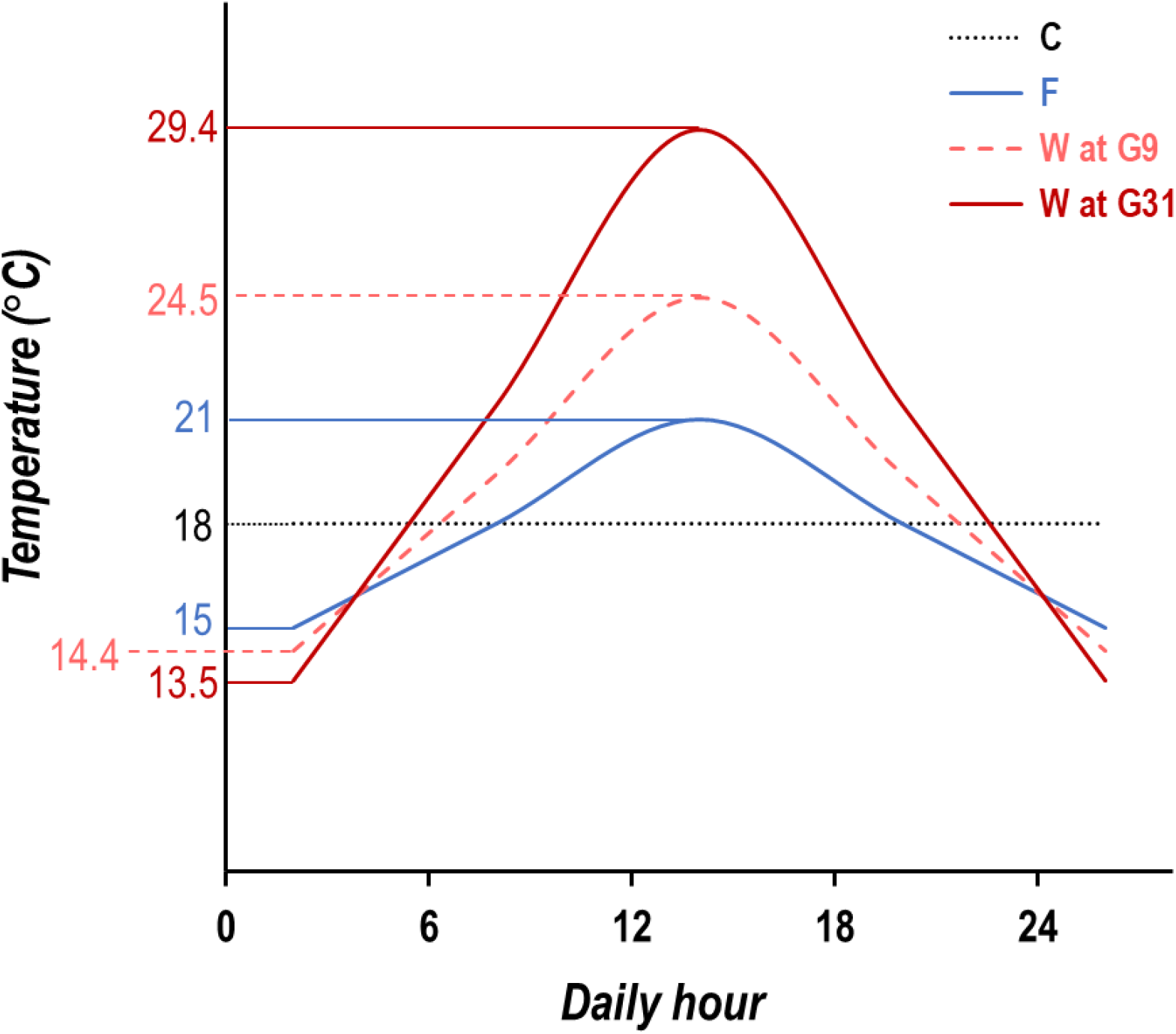
Daily temperature profiles of the experimental regimes after nine (G9) and thirty-one (G31) generations of thermal selection. The warming profile was kept unchanged from generation 20 on.

The PT and NL populations have been kept at 18ºC since their foundation and were already adapted to lab conditions when the two new thermal regimes were created, serving, thus, as controls. As they are expected to be at evolutionary equilibrium, they reflect the ancestral state prior to the start of the F and W selection regimes. All experimental populations were, otherwise, kept under the same conditions, as explained above. Census sizes were, in general, high for all thermal selection regimes with the exceptions referred below (see Figure S1).

The experimental populations were synchronously assayed twice: (1) after 9 generations of thermal evolution (F and W), when the control populations (C) had 79 generations of lab adaptation; and (2) after 31 generations of W selection and 29 generations in the F regime, together with the controls (evolved for 99 generations in the standard lab conditions).

After 9 generations in the warming regime, the experimental populations had been subjected to temperatures ranging from 14.4ºC to 24.5ºC, with a daily average of 19.4ºC (Figure 1). By generation 22, the thermal mean and amplitude increases in the warming regime (peak temperature of 30.2ºC) had to be halted. Particularly low egg-to-adult viability led to significant declines in population size mainly between generations 22 and 24 (see Figure S1). Since then, the warming populations have been kept in the temperature cycle of generation 20, *i*.*e*., with a mean temperature of 21.4ºC, a lower extreme of 13.5ºC, and an upper extreme of 29.4ºC (Figure 1). The temperature increase caused a progressively shorter development time (*i*.*e*., from egg to adult) and, by generation 31, had led to a 96h reduction in the life-cycle length, that became 24 days.

### Thermal plasticity assays

To study the evolution of thermal plasticity, we tested the effect of different thermal environments on the reproduction of adult flies. We analyzed the reproductive performance of eighteen experimental populations (three thermal selection regimes – W, F and C, two histories – NL and PT, and three replicate populations) subjected to three thermal treatments (14ºC, 18ºC, and 24ºC) in two different time points – after nine and 31 generations of thermal evolution; these assays are thereon referred as G9 and G31, respectively. The results for the G9 assay were previously reported in (Santos et al. 2021b). Experimental protocols for both G9 and G31 assays were identical. Briefly, the temperature treatments involved a lifelong exposure to a constant temperature: colder (14ºC), intermediate (18ºC), or warmer (24ºC). These temperatures were chosen considering the range of viable developmental temperatures for *D. subobscura* (6 – 26ºC, (Moreteau et al. 1997; David et al. 2005; Schou et al. 2016)). Maternal effects were minimized by rearing all populations in a common-garden environment, under control conditions (18ºC and 28-day life cycle), for one full generation, before each assay.

In each generation, sixteen pairs of virgin males and females, per population and treatment, were individually assayed with a total of 864 couples. The mating pairs were transferred to fresh medium every other day, for eight days, and the presence of eggs was checked daily. The eggs laid between days six and eight were counted under a stereoscope. Four life history traits were assessed: (1) *age of first reproduction* (number of days since emergence until the first laid egg), where lower values show faster maturity; (2) *fecundity* (total number of eggs laid between days six and eight), which addresses the females reproductive performance when the selective pressures are higher, *i*.*e*., near the age of egg collection for the following generation; (3) *productivity* (number of offspring obtained from the eggs of day eight), which conveys the females’ ability to produce viable progeny; and (4) *juvenile viability* (ratio between productivity and fecundity at day eight), which measures developmental success. To obtain more reliable estimates of viability, only vials with at least five eggs were considered for this trait, with ∼9% of the total number of vials being excluded from this analysis. This was done to adjust for the increase in fungal infections observed in vials with a very low number of eggs.

### Statistical methods

Raw data for the analyses is the mean value for each replicate population and temperature combination (*e*.*g*., three data values for FPT populations in each of the three temperature treatments). Data was analyzed by linear mixed effects models fitted with REML (restricted maximum likelihood). Three factor ANOVAs (Type III Wald F tests, Kenward-Roger degrees of freedom) were done to estimate significance levels of differences between thermal selection regimes, histories and temperature treatments, as well as their interactions. Homoscedasticity and normality were checked. Considering the robustness of the analysis of variance, minor violations were accepted (Knief and Forstmeier 2021). Arcsine transformation was applied to the juvenile viability data. All statistical analyses were performed in R v4.0.4, using the *lme4* (Bates et al. 2015), *car* (Fox and Weisberg 2019), *lawstat* (Hui et al. 2008), *emmeans and ggplot2* (Wickham 2016) packages. Two general models were used to analyze the evolution of thermal reaction norms:

1. *Y = μ + History + Temp + Selection + Gen + AP{History} + History × Temp + History × Selection + Selection × Temp + History × Gen + Temp × Gen + Selection × Gen + Selection × History × Temp + History × Temp × Gen + History × Selection × Gen + Selection × Temp × Gen + Selection × History × Temp × Gen + ε*
2. *Y = μ + History + Temp + Selection + Gen + Block + History × Temp + History × Selection + Selection × Temp + History × Gen + Temp × Gen + Selection × Gen + Selection × History × Temp + History × Temp × Gen + History × Selection × Gen + Selection × Temp × Gen + Selection × History × Temp × Gen + ε*

with *Y* being the trait in study (age of first reproduction, fecundity, productivity, or viability); *History*, as the fixed factor corresponding to geographical origin (with PT and NL as categories); *Selection* as the fixed factor representing the thermal selection regimes (with three categories: Control, Warming, and Fluctuating); *Temp* as the fixed factor corresponding to the three temperature treatments; and *Gen* as the fixed factor with two categories (generations 9 and 31). In these models, interactions between Selection or History and Temperature test for differences in thermal plasticity due to those factors, while interactions between Generation and Temperature indicate temporal changes in plasticity between the two generations. In model (1), *AP{History}* is the random effect consisting of the ancestral replicate population (*i*.*e*., PT_1-3_; NL_1-3_) nested in *History*, from which the replicate populations of the three thermal selection regimes were originated *(e*.*g*., Ancestral PT_1_ generated Control PT_1_, Fluctuating FPT_1_ and Warming WPT_1_). In model (2), *Block* was defined as the random effect, corresponding to the set of same-numbered replicate populations from all thermal regimes, assayed in the same pseudo-randomized experimental rack. For this analysis, we used as raw data in each generation the difference between the mean of each replicate population and the average of the three synchronously assayed control populations (*e*.*g*., FPT_1_-PT, with PT as average of PT_1-3_). As the control populations are long established in the laboratory and, thus, expectedly near evolutionary equilibrium, this procedure allows to remove environmental effects due to the between generation assay asynchrony (*e*.*g*., see (Fragata et al. 2016). In fact, analysis for the control raw data did not show significant variation between generations. Thus, the temporal analyses included standardized data for Warming and Fluctuating populations. Models including or excluding interactions with the random factors (Block or Ancestral Population) were tested and AIC was used to define the best model for each trait. For age of first reproduction and productivity, the model with Block as random effect without its interactions was used; the best suited model for fecundity and viability was the one with Ancestral Population as random effect without its interactions. For viability data analyses, both models (1) and (2) were tested including fecundity of day eight (F8) as a covariate (with models including and excluding its interactions with fixed factors), to account for variation in fecundity across the several thermal treatments. This analysis was also done separately for each selection regime (W and F).

We addressed the evolutionary dynamics of thermal reaction norms by comparing longer (by generation 31) and shorter-term evolution data (by generation 9, published in (Santos et al. 2021b)). We analyzed differences in experimental populations associated with the thermal selective regime imposed (*Selection*, W *vs*. F), the geographical origin (*History*, PT *vs*. NL), the temperature treatments (*Temp*, 14º, 18ºC, and 24ºC), and the two assayed generations (*Gen*, 9 and 31).

Thermal reaction norms were also analyzed after 31 generations of thermal evolution (*i*.*e*., using only data from generation 31). The statistical models used were similar to (1) and (2), but without *Generation* as factor. Raw data for these analyses was the mean absolute value for each replicate population and temperature combination.

Models including and excluding interactions with random effects were assessed with AIC. Considering the traits A1R, fecundity and productivity the best model was model (1) for fecundity and productivity, excluding interactions with the random effect AP; model (2) for age of first reproduction, excluding interactions with the random effect block. For this trait, the model with the lowest AIC was the one defining the random effect AP and F8 as covariate without interactions with fixed or random factors.

## Results

### Evolutionary dynamics of thermal reaction norms

To address the dynamics of temporal changes in thermal reaction norms between generation 31 and 9, we compared the population’s reaction norm after 31 generations of selection considering the three lifelong temperature treatments (14ºC, 18ºC, and 24ºC), with the data after 9 generations (see (Santos et al. 2021b) and Figures 2 and 3).

**Figure 2.**
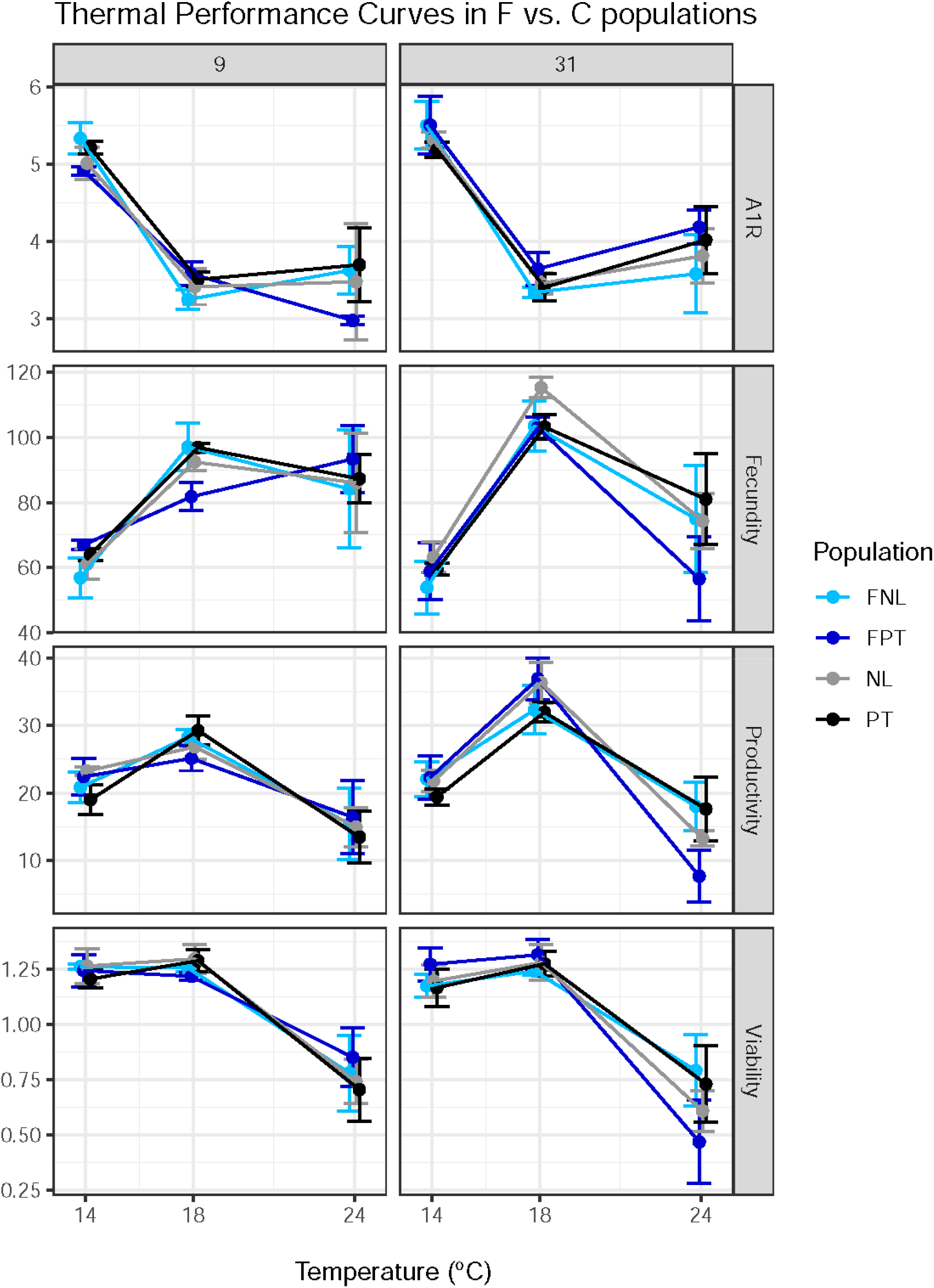
Evolution of the thermal reaction norms of the Fluctuating (F) and Control (C) thermal selection regimes after 9 and 31 generations of selection. Data show the average and 95% confidence intervals for each thermal regime at each time point, with average values of each replicate population as raw data. Viability data is arcsine transformed. A1R – Age of first reproduction.

**Figure 3.**
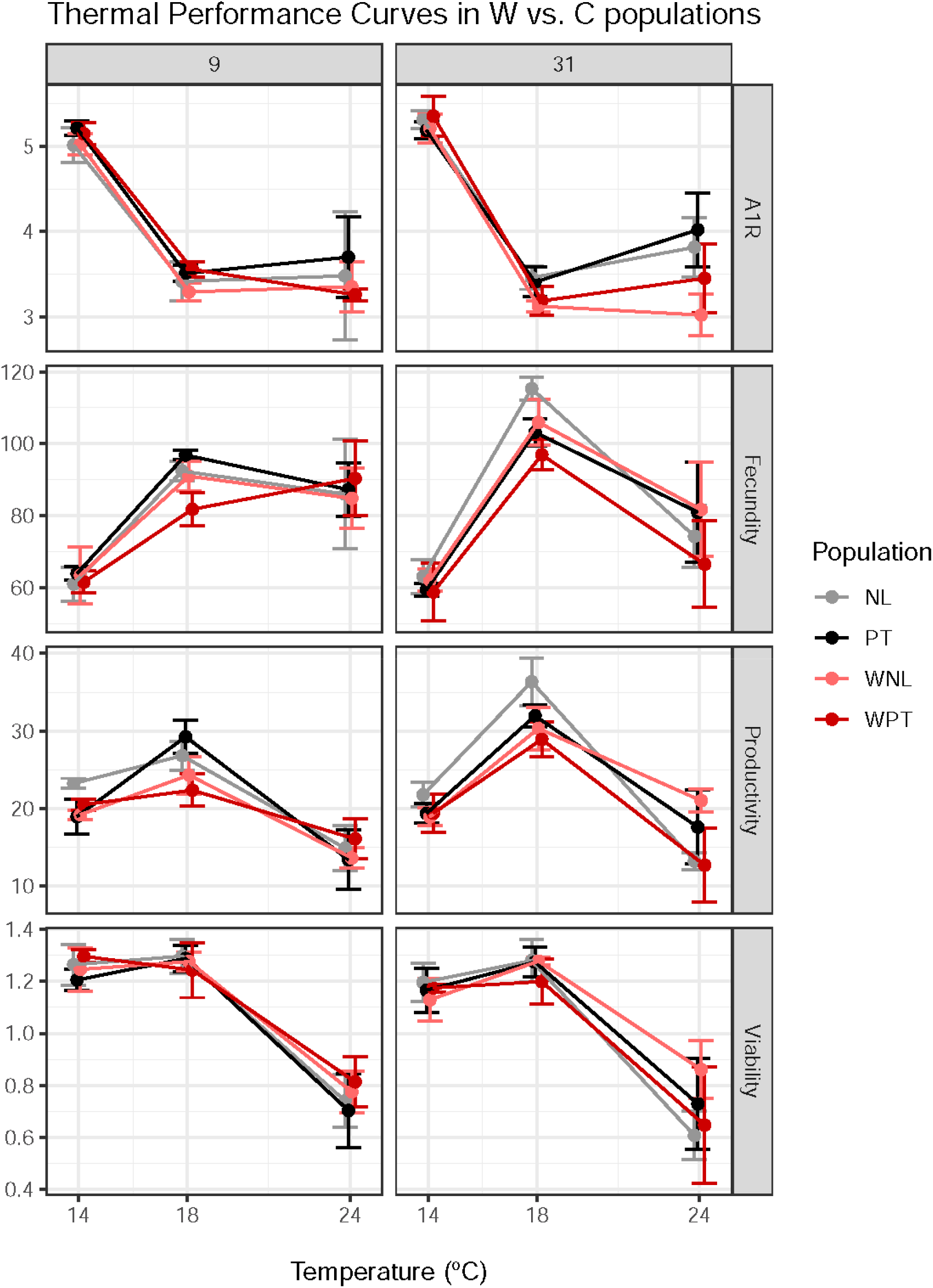
Evolution of the thermal reaction norms of the Warming (W) and Control (C) thermal selection regimes after 9 and 31 generations of selection. Data show the average and 95% confidence intervals for each thermal regime at each time point, with average values of each replicate population as raw data. Viability data is arcsine transformed. A1R – Age of first reproduction.

First, we analyzed the combined effects of thermal selection (W *vs*. F), biogeographical history and their interaction on the reaction norms’ evolution for the three lifelong temperatures in the two assayed generations (Table 1). No overall effects of selection (*Selection x Gen*) or its effects on the reaction norms (*Selection x Temp x Gen*) across generations were observed. The only exception was a significant across-generation difference in age of first reproduction between W and F lines, with an increased performance at 18ºC and 24ºC in the warming populations (see Table 1, A1R, *Selection x Generation*; and Figures 2 and 3). Overall temporal changes between populations with contrasting history were observed for age of first reproduction and viability (see Table 1, *History x Gen*). This effect resulted mainly from the temporal pattern (better performance) of the northern populations in both selection regimes (FNL_1-3_ and WNL_1-3_) compared to the southern populations (see Figures 2 and 3). Changes in reaction norms associated with biogeographical history were found for productivity and viability (see Table 1, *History x Temp*); however, the patterns of variation differed between generations, leading to a significant *History x Temp x Gen* interaction term (see below). Hence, the interpretation of this *History x Temp* interaction should be cautious. In terms of the thermal reaction norm’s evolution, all traits but age of first reproduction showed significant temporal changes due to different geographical histories (see Table 1, *History x Temp x Gen*). This variation was mostly due to the relative performance of southern and northern populations at higher temperatures and across generations (see Figures 2 and 3). After 31 generations of thermal selection, the southern populations showed a general drop in performance at 24ºC, when compared to generation 9, while the northern populations not only maintain their performance but also had an increase in productivity. Given the effect of geographical history on the evolution of thermal reaction norms, additional temporal analyses were performed on each set of populations from the same geographical origin (*e*.*g*., WNL and FNL), see Figures S2 and S3. Temporal changes in thermal reaction norms were significant for productivity in both sets of geographical populations (*Temp x Gen*; northern: F_2,22_=5.64, p<0.011; southern: F_2,22_=8.66, p<0.002), and for fecundity in the southern populations (*Temp x Gen*; southern: F_2,22_=5.54, p<0.012; northern: F_2,22_=1.27, p>0.29). However, no significant temporal changes in reaction norms of populations from different thermal selective regimes were found for any trait (*Selection x Temp x Gen*; p>0.05 in all analyses). Finally, there were no temporal changes in the reaction norms for viability either for the northern or southern populations (*Temp x Gen*; p>0.05).

**Table 1.** Evolutionary dynamics of W and F populations. Data refers to average population values standardized to the controls. The F statistic shows the degrees of freedom of effect and error. Significance levels for F values: p> 0.05 n.s.; 0.05>p>0.01*; 0.01>p>0.001**; p<0.001 ***. Only the model parameters with a significant effect in at least one trait are presented.

| Model parameters | Age of First<br>Reproduction (A1R) | Fecundity | Productivity | Viability |
| --- | --- | --- | --- | --- |
| History*Temp | $F_{2,4} = 2.300$ n.s. | $F_{2,44} = 1.111$ n.s. | $F_{2,46} = 7.250$ ** | $F_{2,43.6} = 3.594$ * |
| History*Gen | $F_{1,28} = 11.08$ ** | $F_{1,44} = 0.007$ n.s. | $F_{1,46} = 2.236$ n.s. | $F_{1,43.1} = 4.613$ * |
| Selection*Gen | $F_{1,28} = 7.185$ * | $F_{1,44} = 0.571$ n.s. | $F_{1,46} = 0.108$ n.s. | $F_{1,43.0} = 0.001$ n.s. |
| History*Temp*Gen | $F_{2,28} = 2.911$ n.s. | $F_{2,44} = 6.054$ ** | $F_{2,46} = 14.811$ *** | $F_{2,43.8} = 5.459$ ** |

Second, we tested the evolutionary change of each thermal regime’s reaction norm between generations 9 and 31. We did not detect a significant effect of the intercept or significant temporal changes in the overall performance of any thermal regime (Table S1, *intercept*, and *Gen*). A significant effect of temperature was observed for age of first reproduction and productivity in the W populations (Table S1, *Temp*) but no temporal changes in the reaction norms were observed for any trait (Table S1, *Gen x Temp*). Also, historical variation did not present a significant effect either on overall performance (Table S1, *History*) or across generations (Table S1, *History x Gen*). However, history had a significant effect on the overall shape of the reaction norms for productivity, in both warming and fluctuating populations, but not for the other traits (Table S1, *History x Temp*). In addition, a significant *History x Temp x Gen* interaction was observed for fecundity in the F populations and productivity in both F and W populations, indicating historical differences in the evolution of thermal reaction norms (see Table S1 and Figures 2 and 3, see also below). This temporal analysis was also done separately for each combination of Selection and History populations (*i*.*e*., WPT, WNL, FPT, and FNL) – see Table S2. Temporal changes in thermal reaction norms (Table S2, *Temp x Gen*) were observed for fecundity and productivity in the FPT populations, and productivity for the WNL populations. Trait changes in FPT populations resulted from the temporal decline in performance at higher temperatures, while a temporal increase in productivity was observed for the WNL populations. This productivity increase occured at both lower (approaching control values) and higher temperatures (diverging from control values) – see Figures 2 and 3.

### Longer-term changes of thermal reaction norms after 31 generations under varying thermal environments

We first analyzed the differences in thermal reaction norms between each selective regime, fluctuating or warming, and the controls (Figures 2 and 3, respectively). Differences between temperature treatments were observed for all traits in both F *vs*. C and W *vs*. C comparisons (significant *Temp* factor, Tables S3 and S4). Different traits showed different thermal responses: both colder (14ºC) and warmer (24ºC) temperatures caused a decline in fecundity and productivity, when compared to control conditions (18ºC); age of first reproduction showed reduced performance at lower but not at warmer temperatures, while viability showed the reverse pattern (lower performance at the warmest temperature) – see Figures 2 and 3; Tables S3 and S4. No overall effects of biogeographical history or thermal selection were observed for most traits. The exception was for age of first reproduction, where W populations showed significantly lower values (see Table S3 and Figure 3).

The reaction norms of populations from different selection regimes (F *vs*. C or W *vs*. C) or from different biogeographical histories (Portugal *vs*. The Netherlands) were also not significantly different (*Selection x Temp* and *History x Temp* interactions, respectively, Tables S3 and S4). A significant triple interaction *History x Selection x Temp* was observed for productivity in the F *vs*. C comparison (Table S4, Figure 2), indicating that populations from distinct geographical origin respond differently to thermal selection. This effect was mainly due to a lower performance at 24ºC of the southern populations subjected to the fluctuating regime (FPT).

Finally, we compared the performance of populations evolving under the two varying thermal environments (fluctuating *vs*. warming). Like what was found above, differences between temperature treatments were observed for all traits (Table S5, *Temp*). Overall differences in performance between selective regimes were only found for age of first reproduction (Table S5, *Selection*) while no differences were found in the reaction norms between F and W regimes for any trait (Table S5, *Selection x Temp*). Interestingly, differences in the reaction norms of populations with diverse biogeographical histories were found for productivity, mostly due to a better performance of northern populations at higher temperatures (Table S5, *History x Temp*).

## Discussion

The temperature increase and fast-changing environmental conditions associated with global warming impose great stress on ectothermic animals and adaptation can be key for their survival. Here, we present a 31 generation-long experimental evolution study addressing the evolutionary dynamics of thermal plasticity in populations of *D. subobscura*. Two populations with distinct biogeographical histories (northern and southern Europe) were subjected to different thermal selection environments (daily fluctuating and global warming scenarios).

### Historical differences lead to contrasting evolutionary dynamics of thermal reaction norms

We found that the evolutionary dynamics of thermal reaction norms across 22 generations of evolution (generation 9 *vs*. 31) for several life history traits varied between populations of contrasting geographical origin. This temporal variation in plasticity was mostly driven by a differential performance at higher temperatures, with the southern populations decreasing their reproductive performance across generations, while their northern counterparts presented the opposite pattern; this effect was more evident in their progeny productivity (Figures 2 and 3, Table 1). Geographical variation in the shape of thermal reaction norms has been found in some studies of thermal response in ectotherms (*e*.*g*., (Trotta et al. 2006; Berger et al. 2013; Austin and Moehring 2019; Klepsatel et al. 2019) but not in others (Klepsatel et al. 2013; Clemson et al. 2016). Particularly in *D. subobscura*, variation in thermal reaction norms has been observed in populations from Sweden and Spain, with northern populations performing worse at higher temperatures (Porcelli et al. 2017). One could, thus, expect northern *D. subobscura* natural populations to exhibit a steeper evolutionary response to rising temperatures, as these populations are likely further away from the new optimum in a warming scenario – and assuming that all populations evolve towards the same adaptive peak (Matos et al. 2002; Losos 2011). Prior to the establishment of the thermal regimes, our northern populations did not show a worse performance at higher temperatures when compared to their southern counterparts (Simões et al. 2020; Santos et al. 2021a). However, after 9 generations of thermal selection, though there was no robust evidence for historical effects, we detected a tendency for lower performance of the northern populations at higher temperatures across traits (Santos et al. 2021b). In this sense, the contrasting temporal dynamics observed between geographical populations at higher temperatures agrees with these expectations. Nevertheless, and assuming convergence, this variation in dynamics would lead to lower historical differences after 31 generations of thermal evolution. On the contrary, we found that populations are diverging through time, with the northern populations increasing their performance at higher temperatures and southern ones decreasing theirs. This pattern is particularly evident for productivity. This is an interesting finding as it suggests that, given enough time, populations historically from cooler climates might better overcome the current thermal challenges. The increased performance in the northern populations could result from a better developmental acclimation to higher temperatures, as both developmental and adult stages were subjected to the same temperature, which is worthy of further investigation. The presence of high genomic differentiation between the southern and northern populations at the start of our experiment might also play a role in the observed patterns. This is corroborated by previous findings in other *D. subobscura* populations that were founded from the same natural locations and still exhibit a high degree of genomic differentiation, even after 50 generations of lab evolution (Seabra et al. 2018). Further studies are needed to address the relevant genomic variation between the evolving populations, so that we understand the driving force behind the geographical variation and the temporal increase in performance in the northern populations.

Our results suggest non-linear and complex evolutionary trajectories in which an interplay between geographical backgrounds and the varying thermal environments has an important role. The absence of clear geographical variation in our short term study (Santos et al. 2021b) and the later emergence of such differences reinforces the need for caution when interpreting and extrapolating short term evolution patterns to longer term evolution.

### Evolution under different thermal selection regimes does not lead to different thermal reaction norms

Considering the higher thermal variation in the warming *vs*. the fluctuating environment with much more pronounced thermal extremes, one could expect a higher performance of the warming-evolved populations at higher and lower temperatures, as well as a lower performance at intermediate ones. This expectation is based on a possible trade-off between thermal breath and maximal performance (Huey and Kingsolver 1993). However, we did not find differences in the evolution of thermal reactions norms between the warming and fluctuating populations, despite their geographical origin. It might be the case that enzymatic constraints limit the evolution of improved performance at both extremes (Angilletta et al. 2003), which, in turn, might hinder the evolution of a more generalist performance of the warming populations. In fact, other experimental evolution studies have seldom verified the evolution of such predicted phenotypes as a response to thermal environmental heterogeneity, including the evolution of thermal specialists *vs*. generalists (*e*.*g*., (Berger et al. 2014; Condon et al. 2014; Manenti et al. 2015) but see (Le Vinh Thuy et al. 2016).

### Can populations adapt to a global warming environment?

We did not find a clear long-term adaptive response to higher temperatures after 31 generations of thermal selection under a warming scenario, despite the trend of improved productivity (*i*.*e*., number of offspring) of the northern populations (see Figure 3 and Tables S2 and S3). Other experimental evolution studies addressing adaptation to cumulative higher temperatures have also failed to find a clear adaptive response to these more stressful environments (*e*.*g*., (Schou et al. 2014; Kinzner et al. 2019). Constrained plastic response of physiological traits at high temperatures was also observed in comparative studies across species (Gunderson and Stillman 2015; Schou et al. 2016; Sørensen et al. 2016; MacLean et al. 2019). Several causes might contribute to the absence of a clear adaptive response, which was particularly evident in our southern populations. Evolutionary constraints, such as trade-offs or an overall low genetic variation for the measured traits, might play an important role (Hoffmann et al. 2017; Kristensen et al. 2020). Methodological issues, such as the decoupling between the assay environments and the selection regimes (*e*.*g*., constant *vs*. fluctuating temperatures, low *vs*. high density reproduction) might also contribute. Another possible explanation is that a part of the population’s response to the thermal conditions might be environmentally-based, resulting, for instance, from epigenetic and/or maternal environmental effects (*e*.*g*., (Hoffmann and Bridle 2021). That being true, such effects would not be detected in our experimental setup as they could have been mitigated or removed by the one-generation common garden protocol at 18ºC imposed prior to the plasticity assays. The impact of common garden procedures and the environment in which they occur is an issue worthy of further consideration and analysis, particularly in the context of experimental evolution.

The selective pressure experienced by populations in the new thermal environment is also an important point to consider. There were times during thermal evolution when the selective pressures in the warming environment were most likely very high, as there were severe drops in the W population sizes’ due to extremely low juvenile viability. Nevertheless, in more recent generations, the populations showed an obvious improvement in their ability to endure the maintenance conditions, with consistently higher census sizes (Figure S1). This makes us wonder why such an apparent adaptive response is not clearly detected in our life-history assays. We believe that this might stem from a decoupling between the test environment and the imposed selection environment. However, even if adaptation to the different thermal conditions occurred, it did not produce conspicuous changes in the thermal reaction norms. In future experiments, it will be important to test the exact environment in which the populations evolve (*i*.*e*., local adaptation assays), alongside with changes in the thermal reaction norms measured at constant temperatures.

### Final remarks

We, here, report differential plasticity during evolution under thermally varying environments, driven by the contrasting biogeographical history of the evolving populations. The northern European populations appear to be coping better with these dynamic thermal environments, as they showed a temporal increase in performance at higher temperatures, particularly in terms of productivity. Contrary to our expectations, the evolutionary changes in thermal reaction norms did not lead to an across-trait increased performance at lower and higher temperatures after long-term thermal selection. Remarkably, our study shows a high impact of the previous biogeographical history in the population’s thermal evolutionary response, even after tens of generations evolving in the same environment. These results highlight the importance of extending the evolutionary studies to populations from different geographical sources, which have adapted to local environmental conditions, to make more accurately intraspecific generalizations. Our findings reinforce the likely complex nature of thermal responses and illustrate the hardships of predicting evolutionary responses to global warming.

## Acknowledgments

The authors thank Inês Fragata for help in the graphic data presentation. This study is financed by Portuguese National Funds through ‘Fundação para a Ciência e a Tecnologia’ (FCT) within the projects PTDC/BIA-EVL/28298/2017 and cE3c Unit FCT funding project UIDB/00329/2020. PS and ASQ are funded by national funds (OE), through FCT, in the scope of the framework contract foreseen in the numbers 4, 5 and 6 of the article 23^rd^, of the Decree-Law 57/2016, of August 29, changed by Law 57/2017, of July 19. MS is funded by grants CGL2017-89160P from Ministerio de Economía y Competitividad (Spain; co-financed with the European Union FEDER funds), and 2017SGR 1379 from Generalitat de Catalunya.

## Notes

### Competing Interest Statement

The authors have declared no competing interest.

