## Supporting information for "Biogeographical history shapes evolution of reproduction in a global warming scenario"

### Supplementary figure legends

**Figure S1.** Census sizes of the experimental populations (estimated on the day of reproduction) throughout the thermal selection protocol for **a)** control, **b)** fluctuating, and **c)** warming populations.

**Figure S2.** Evolution of the thermal reaction norms of the Southern populations (FPT and WPT) after 9 and 31 generations of selection. Data show the average and 95% confidence intervals for each thermal regime at each time point, with average values of each replicate population as raw data. A1R – Age of first reproduction.

**Figure S3.** Evolution of the thermal reaction norms of the Northern populations (FNL and WNL) after 9 and 31 generations of selection. Data show the average and 95% confidence intervals for each thermal regime at each time point, with average values of each replicate population as raw data. A1R – Age of first reproduction.

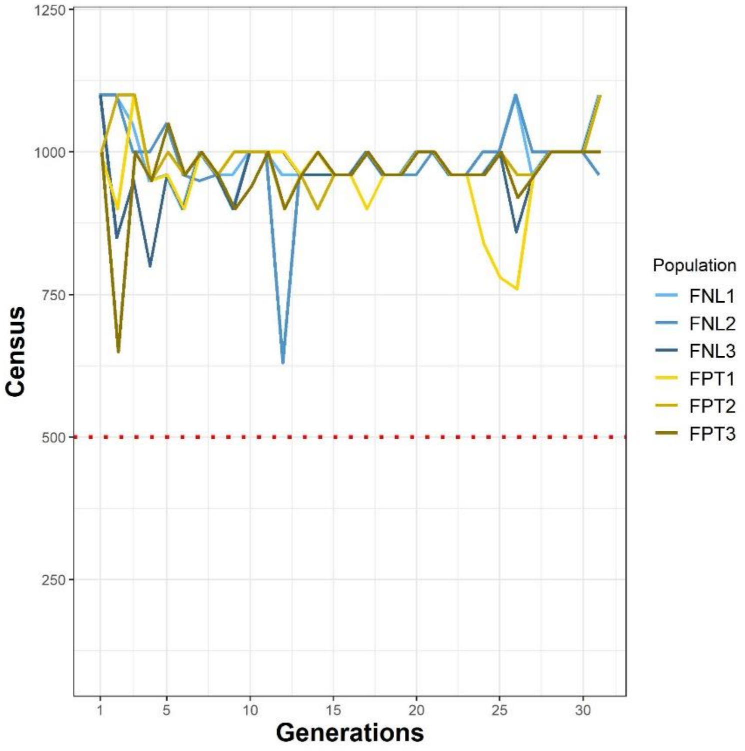


Figure S1a)

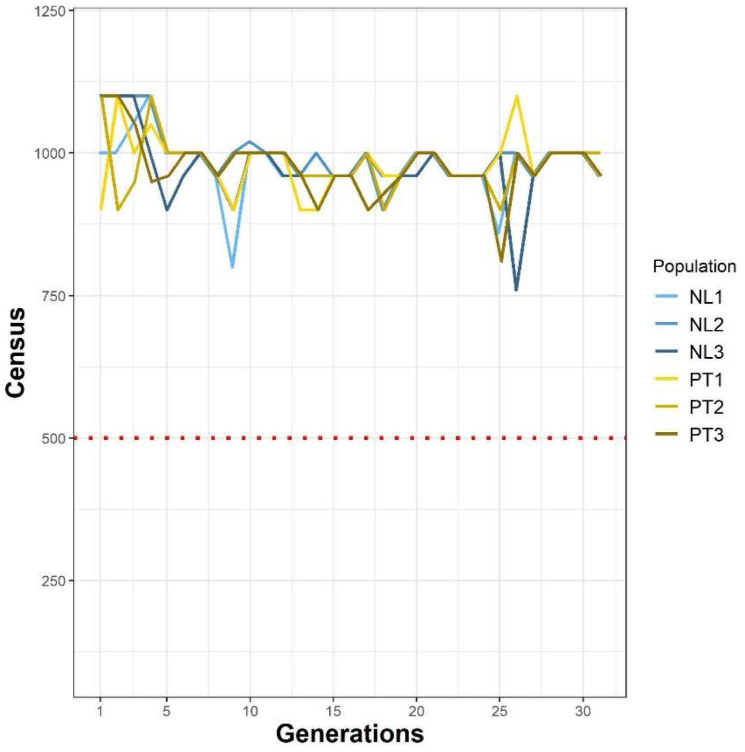


Figure S1b)

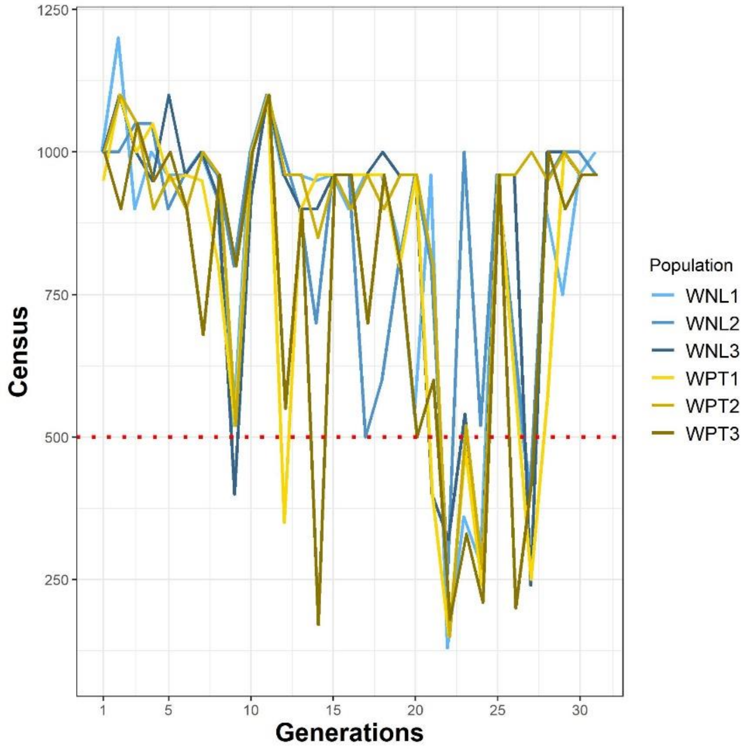
1250
1000
750
500
250
Census
1
5
10
15
20
25
30
Generations
Population
WNL1
WNL2
WNL3
WPT1
WPT2
WPT3

Figure S1c)

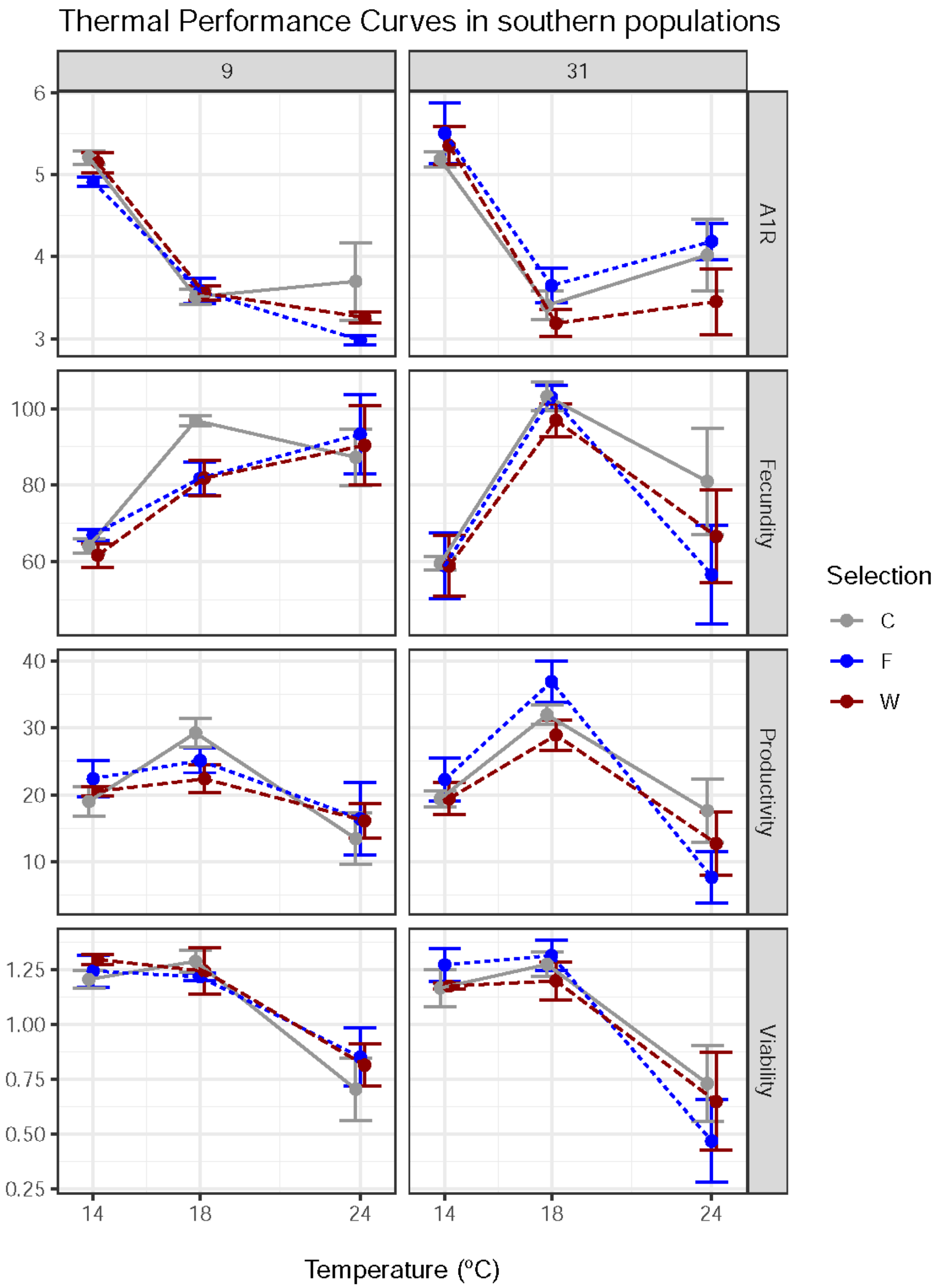
Thermal Performance Curves in southern populations
9
31
A1R
Fecundity
Productivity
Viability
6
5
4
3
100
80
60
40
30
20
10
1.25
1.00
0.75
0.50
0.25
14
18
24
Temperature (ºC)
Selection
C
F
W

Figure S2

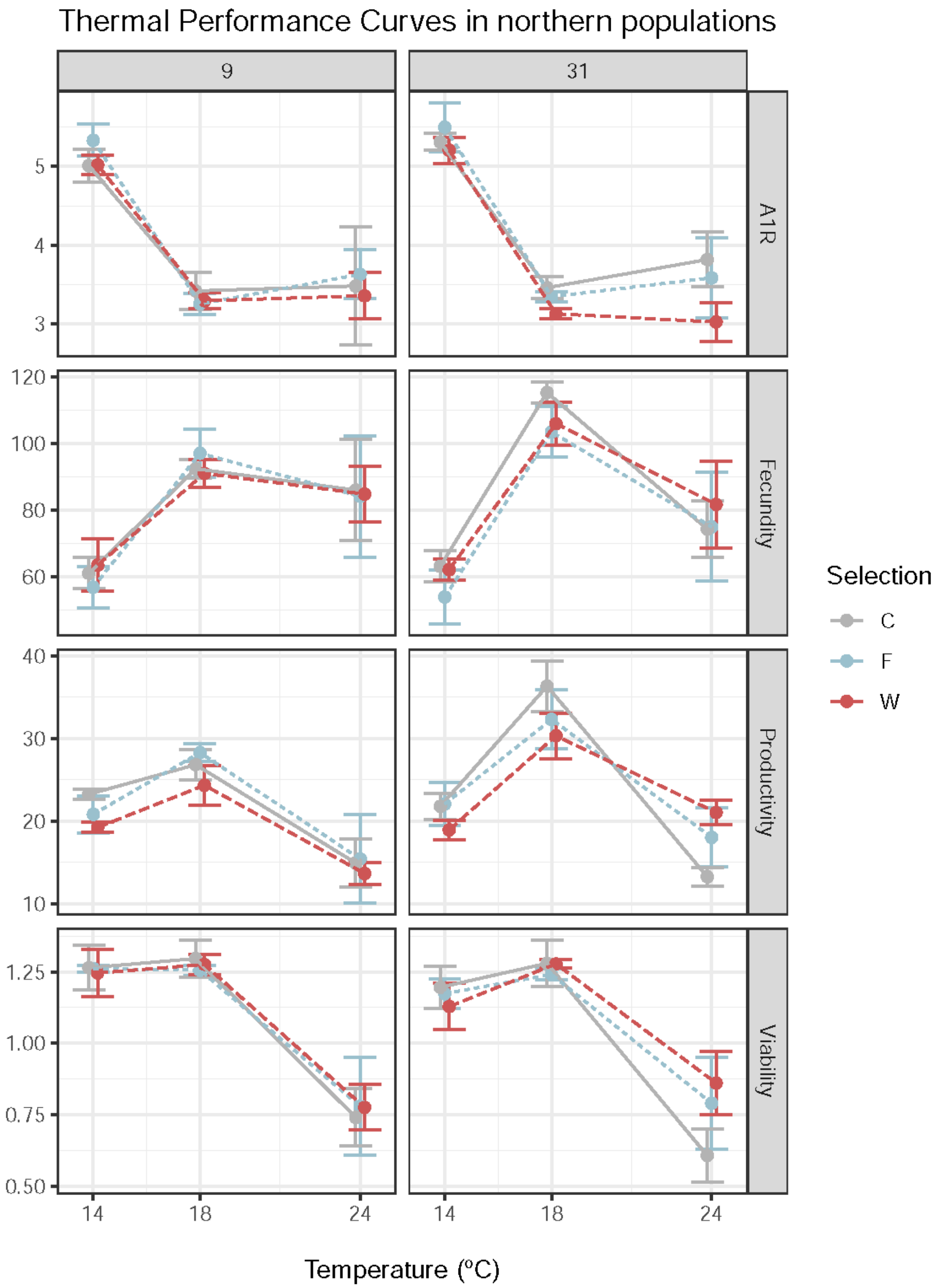

Thermal Performance Curves in northern populations
9
31
A1R
Fecundity
Productivity
Viability
5
4
3
120
100
80
60
40
30
20
10
1.25
1.00
0.75
0.50
14
18
24
14
18
24
Temperature (ºC)
Selection
C
F
W


Figure S3

### Supplementary tables

**Table S1.** Temporal changes in plasticity for each thermal selection regime (G9 *vs.* G31): **a)** Controls, **b)** Warming, and **c)** Fluctuating populations. In a) data refers to the average population values, in b) and c) data refers to average population values standardized to the controls. The F statistic shows the degrees of freedom of effect and error. Significance levels for F values: p> 0.05 n.s.; 0.05>p>0.01*; 0.01>p>0.001**; p<0.001 ***.

a)

| Trait | $F_{(df1, df2)}$ - Intercept | $F_{(df1, df2)}$ - Gen | $F_{(df1, df2)}$ - History | $F_{(df1, df2)}$ - Temp | $F_{(df1, df2)}$ - Temp x Gen | $F_{(df1, df2)}$ - History x Gen | $F_{(df1, df2)}$ - History x Temp | $F_{(df1, df2)}$ - History x Temp x Gen |
|---|---|---|---|---|---|---|---|---|
| Age of First Reproduction (A1R) | $F_{1,20}$ = 0.634 n.s. | $F_{1,20}$ = 0.634 n.s. | $F_{1,4}$ = 0.176 n.s. | $F_{2,20}$ = 34.229 *** | $F_{2,20}$ = 0.323 n.s. | $F_{1,20}$ = 0.195 n.s. | $F_{2,20}$ = 0.114 n.s. | $F_{2,20}$ = 0.060 n.s. |
| Fecundity (F6-8) | $F_{1,20}$ = 0.122 n.s. | $F_{1,20}$ = 0.122 n.s. | $F_{1,4}$ = 0.0003 n.s. | $F_{2,20}$ = 30.469 *** | $F_{2,20}$ = 2.752 n.s. | $F_{1,20}$ = 0.501 n.s. | $F_{2,20}$ = 0.289 n.s. | $F_{2,20}$ = 0.576 n.s. |
| Productivity | $F_{1,20}$ = 2.464 n.s. | $F_{1,20}$ = 2.464 n.s. | $F_{1,4}$ = 0.429 n.s. | $F_{2,20}$ = 42.898 *** | $F_{2,20}$ = 1.867 n.s. | $F_{1,20}$ = 0.010 n.s. | $F_{2,20}$ = 0.904 n.s. | $F_{2,20}$ = 1.657 n.s. |
| Viability | $F_{1,22.4}$ = 0.540 n.s. | $F_{1,22.4}$ = 0.540 n.s. | $F_{1,3.5}$ = 0.0003 n.s. | $F_{2,21.2}$ = 41.693 *** | $F_{2,20.4}$ = 0.025 n.s. | $F_{1,19.7}$ = 0.371 n.s. | $F_{2,19.9}$ = 0.133 n.s. | $F_{2,20.4}$ = 0.131 n.s. |

b)

| Trait | $F_{(df1, df2)}$ - Intercept | $F_{(df1, df2)}$ - Gen | *$F_{(df1, df2)}$ - History* | $F_{(df1, df2)}$ - Temp | $F_{(df1, df2)}$ - Temp x Gen | $F_{(df1, df2)}$ - History x Gen | $F_{(df1, df2)}$ - History x Temp | $F_{(df1, df2)}$ - History x Temp x Gen |
|---|---|---|---|---|---|---|---|---|
| Age of First Reproduction (A1R) | $F_{1,4}$ = 6.078 n.s. | $F_{1,20}$ = 3.634 n.s. | $F_{1,4}$ = 0.136 n.s. | $F_{2,20}$ = 7.993 ** | $F_{2,20}$ = 1.738 n.s. | $F_{1,20}$ = 1.839 n.s. | $F_{2,20}$ = 0.347 n.s. | $F_{2,20}$ = 0.744 n.s. |
| Fecundity (F6-8) | $F_{1,4}$ = 1.618 n.s. | $F_{1,20}$ = 0.184 n.s. | $F_{1,4}$ = 1.017 n.s. | $F_{2,20}$ = 1.161 n.s. | $F_{2,20}$ = 0.160 n.s. | $F_{1,20}$ = 0.018 n.s. | $F_{2,20}$ = 0.175 n.s. | $F_{2,20}$ = 2.124 n.s. |
| Productivity | $F_{1,4}$ = 6.313 n.s. | $F_{1,20}$ = 0.010 n.s. | $F_{1,4}$ = 0.114 n.s. | $F_{2,20}$ = 5.546 * | $F_{2,20}$ = 0.011 n.s. | $F_{1,20}$ = 2.581 n.s. | $F_{2,20}$ = 3.972 * | $F_{2,20}$ = 7.622 ** |
| Viability | $F_{1,5.2}$ = 0.013 n.s. | $F_{1,19.0}$ = 0.406 n.s. | $F_{1,3.9}$ = 0.319 n.s. | $F_{2,19.1}$ = 1.719 n.s. | $F_{2,19.0}$ = 0.029 n.s. | $F_{1,19.0}$ = 3.606 n.s. | $F_{2,19.1}$ = 3.098 n.s. | $F_{2,19.1}$ = 2.955 n.s. |

c)

| Trait | $F_{(df1, df2)}$ - Intercept | $F_{(df1, df2)}$ - Gen | $F_{(df1, df2)}$ - History | $F_{(df1, df2)}$ - Temp | $F_{(df1, df2)}$ - Temp x Gen | $F_{(df1, df2)}$ - History x Gen | $F_{(df1, df2)}$ - History x Temp | $F_{(df1, df2)}$ - History x Temp x Gen |
|---|---|---|---|---|---|---|---|---|
| Age of First Reproduction (A1R) | $F_{1,4}$ = 0.002 n.s. | $F_{1,20}$ = 2.652 n.s. | $F_{1,4}$ = 0.075 n.s. | $F_{2,20}$ = 1.812 n.s. | $F_{2,20}$ = 0.139 n.s. | $F_{1,20}$ = 8.044 * | $F_{2,20}$ = 2.114 n.s. | $F_{2,20}$ = 1.753 n.s. |
| Fecundity (F6-8) | $F_{1,4}$ = 0.966 n.s. | $F_{1,20}$ = 1.951 n.s. | $F_{1,4}$ = 0.026 n.s. | $F_{2,20}$ = 0.146 n.s. | $F_{2,20}$ = 0.792 n.s. | $F_{1,20}$ = 0.0002 n.s. | $F_{2,20}$ = 1.054 n.s. | $F_{2,20}$ = 3.795 * |
| Productivity | $F_{1,4}$ = 0.000 n.s. | $F_{1,20}$ = 0.122 n.s. | $F_{1,4}$ = 0.006 n.s. | $F_{2,20}$ = 0.396 n.s. | $F_{2,20}$ = 1.539 n.s. | $F_{1,20}$ = 0.453 n.s. | $F_{2,20}$ = 3.884 * | $F_{2,20}$ = 8.015 ** |
| Viability | $F_{1,3.8}$ = 0.420 n.s. | $F_{1,20.0}$ = 0.005 n.s. | $F_{1,3.5}$ = 0.071 n.s. | $F_{2,20.8}$ = 0.732 n.s. | $F_{2,20.9}$ = 0.176 n.s. | $F_{1,19.4}$ = 1.919 n.s. | $F_{2,20.3}$ = 0.962 n.s. | $F_{2,21.1}$ = 2.436 n.s. |

**Table S2.** Temporal changes in plasticity for each combination or thermal selection and history regimes (G9 *vs.* G31): **a)** FNL, **b)** FPT, **c)** WNL, and **d)** WPT. Data refers to average population values standardized to the controls. The F statistic shows the degrees of freedom of effect and error. Significance levels for F values: p> 0.05 n.s.; 0.05>p>0.01*; 0.01>p>0.001**; p<0.001 ***.

a) FNL

| Trait | $F_{(df1, df2)}$ - Gen | $F_{(df1, df2)}$ - Temp | $F_{(df1, df2)}$ - Temp x Gen |
|---|---|---|---|
| Age of First Reproduction (A1R) | $F_{1,10}$ = 0.680 n.s. | $F_{2,10}$ = 1.693 n.s. | $F_{2,10}$ = 0.448 n.s. |
| Fecundity (F6-8) | $F_{1,10}$ = 1.093 n.s. | $F_{2,10}$ = 0.256 n.s. | $F_{2,10}$ = 0.670 n.s. |
| Productivity | $F_{1,10}$ = 0.083 n.s. | $F_{2,10}$ = 1.879 n.s. | $F_{2,10}$ = 2.604 n.s. |
| Viability | $F_{1,10.8}$ = 1.034 n.s. | $F_{2,9.4}$ = 1.922 n.s. | $F_{2,10.2}$ = 0.843 n.s. |

b) FPT

| Trait | $F_{(df1, df2)}$ - Gen | $F_{(df1, df2)}$ - Temp | $F_{(df1, df2)}$ - Temp x Gen |
|---|---|---|---|
| Age of First Reproduction (A1R) | $F_{1,10}$ = 10.756 ** | $F_{2,10}$ = 2.276 n.s. | $F_{2,10}$ = 1.522 n.s. |
| Fecundity (F6-8) | $F_{1,10}$ = 1.021 n.s. | $F_{2,10}$ = 1.032 n.s. | $F_{2,10}$ = 4.244 * |
| Productivity | $F_{1,10}$ = 0.380 n.s. | $F_{2,10}$ = 2.260 n.s. | $F_{2,10}$ = 5.774 * |
| Viability | $F_{1,9.2}$ = 0.970 n.s. | $F_{2,10.1}$ = 0.346 n.s. | $F_{2,10.2}$ = 1.253 n.s. |

c) WNL

| Trait | $F_{(df1, df2)}$ - Gen | $F_{(df1, df2)}$ - Temp | $F_{(df1, df2)}$ - Temp x Gen |
|---|---|---|---|
| Age of First Reproduction (A1R) | $F_{1,10}$ = 5.403 * | $F_{2,10}$ = 2.801 n.s. | $F_{2,10}$ = 1.436 n.s. |
| Fecundity (F6-8) | $F_{1,10}$ = 0.052 n.s. | $F_{2,10}$ = 0.727 n.s. | $F_{2,10}$ = 0.671 n.s. |
| Productivity | $F_{1,10}$ = 2.450 n.s. | $F_{2,10}$ = 10.942 ** | $F_{2,10}$ = 6.536 * |
| Viability | $F_{1,9.0}$ = 0.941 n.s. | $F_{2,9.0}$ = 5.681 * | $F_{2,9.1}$ = 2.644 n.s. |

d) WPT

| Trait | $F_{(df1, df2)}$ - Gen | $F_{(df1, df2)}$ - Temp | $F_{(df1, df2)}$ - Temp x Gen |
|---|---|---|---|
| Age of First Reproduction (A1R) | $F_{1,10}$ = 0.149 n.s. | $F_{2,10}$ = 5.498 * | $F_{2,10}$ = 1.051 n.s. |
| Fecundity (F6-8) | $F_{1,10}$ = 0.148 n.s. | $F_{2,10}$ = 0.682 n.s. | $F_{2,10}$ = 1.613 n.s. |
| Productivity | $F_{1,10}$ = 0.808 n.s. | $F_{2,10}$ = 2.148 n.s. | $F_{2,10}$ = 2.668 n.s. |
| Viability | $F_{1,9.0}$ = 2.549 n.s. | $F_{2,9.1}$ = 0.672 n.s. | $F_{2,9.1}$ = 0.555 n.s. |

**Table S3.** Thermal reaction norms of the W and C populations, after 31 generations of thermal selection: **a)** ANOVA table and **b)** Tukey HSD tests for the significant factor *Temp*. Data refers to average population values. The F statistic shows the degrees of freedom of effect and error. Significance levels for F values: p> 0.05 n.s.; 0.05>p>0.01*; 0.01>p>0.001**; p<0.001 ***.

a)

| 10 | Model parameters | $F_{(df1, df2)}$ |
|---|---|---|
| Age of First Reproduction (A1R) | History | $F_{1,22}$ = 0.623 n.s. |
| | Selection | $F_{1,22}$ = 4.801 * |
| | Temp | $F_{2,22}$ = 76.140 *** |
| | History*Selection | $F_{1,22}$ = 0.519 n.s. |
| | History*Temp | $F_{2,22}$ = 0.537 n.s. |
| | Selection*Temp | $F_{2,22}$ = 2.145 n.s. |
| | History*Selection*Temp | $F_{2,22}$ = 0.027 n.s. |
| Fecundity | History | $F_{1,4}$ = 1.749 n.s. |
| | Selection | $F_{1,20}$ = 0.766 n.s. |
| | Temp | $F_{2,20}$ = 31.998 *** |
| | History*Selection | $F_{1,20}$ = 0.440 n.s. |
| | History*Temp | $F_{2,20}$ = 0.237 n.s. |
| | Selection*Temp | $F_{2,20}$ = 0.194 n.s. |
| | History*Selection*Temp | $F_{2,20}$ = 0.730 n.s. |
| Productivity | History | $F_{1,4}$ = 1.617 n.s. |
| | Selection | $F_{1,20}$ = 0.961 n.s. |
| | Temp | $F_{2,20}$ = 38.662 *** |
| | History*Selection | $F_{1,20}$ = 0.556 n.s. |
| | History*Temp | $F_{2,20}$ = 0.134 n.s. |
| | Selection*Temp | $F_{2,20}$ = 1.272 n.s. |
| | History*Selection*Temp | *$F_{2,20}$ = 2.907 m.s.* |
| Viability | F8 | $F_{1,21}$ = 1.665 n.s. |
| | History | $F_{1,4}$ = 0.625 n.s. |
| | Selection | $F_{1,20}$ = 0.163 n.s. |
| | Temp | $F_{2,21}$ = 30.628 *** |
| | History*Selection | $F_{1,19}$ = 0.896 n.s. |
| | History*Temp | $F_{2,19}$ = 0.056 n.s. |
| | Selection*Temp | $F_{2,19}$ = 0.359 n.s. |
| | History*Selection*Temp | $F_{2,20}$ = 1.532 n.s. |

b)

| Trait | Model parameters | $F_{(df1, df2)}$ |
|---|---|---|
| Age of First Reproduction (A1R) | Temp | $F_{2,22}$ = 76.140 *** |
| | 14 vs 18 | t.ratio = 11.411 *** |
| | 14 vs 24 | t.ratio = 9.774 *** |
| | 18 vs 24 | t.ratio = - 1.636 n.s. |
| Fecundity (F6-8) | Temp | $F_{2,20}$ = 31.998 *** |
| | 14 vs 18 | t.ratio = - 7.862 *** |
| | 14 vs 24 | t.ratio = - 2.653 * |
| | 18 vs 24 | t.ratio = 5.210 *** |
| Productivity | Temp | $F_{2,20}$ = 38.662 *** |
| | 14 vs 18 | t.ratio = - 6.420 *** |
| | 14 vs 24 | t.ratio = 1.993 n.s. |
| | 18 vs 24 | t.ratio = 8.414 *** |
| Viability | Temp | $F_{2,21}$ = 30.628 *** |
| | 14 vs 18 | t.ratio = - 1.784 n.s. |
| | 14 vs 24 | t.ratio = 3.891 ** |
| | 18 vs 24 | t.ratio = 7.455 *** |

**Table S4.** Thermal reaction norms of the F and C populations, after 31 generations of thermal selection: **a)** ANOVA table and **b)** Tukey HSD tests for the significant factor *Temp*. Data refers to average population values. The F statistic shows the degrees of freedom of effect and error. Significance levels for F values: p> 0.05 n.s.; 0.05>p>0.01*; 0.01>p>0.001**; p<0.001 ***.

a)

| Trait | Model parameters | $F_{(df1, df2)}$ |
|---|---|---|
| Age of First Reproduction (A1R) | History | $F_{1,22}$ = 0.914 n.s. |
| | Selection | $F_{1,22}$ = 0.331 n.s. |
| | Temp | $F_{2,22}$ = 49.707 *** |
| | History*Selection | $F_{1,22}$ = 0.805 n.s. |
| | History*Temp | $F_{2,22}$ = 0.678 n.s. |
| | Selection*Temp | $F_{2,22}$ = 0.263 n.s. |
| | History*Selection*Temp | $F_{2,22}$ = 0.065 n.s. |
| Fecundity | History | $F_{1,4}$ = 0.457 n.s. |
| | Selection | $F_{1,20}$ = 2.285 n.s. |
| | Temp | $F_{2,20}$ = 31.417 *** |
| | History*Selection | $F_{1,20}$ = 0.027 n.s. |
| | History*Temp | $F_{2,20}$ = 0.201 n.s. |
| | Selection*Temp | $F_{2,20}$ = 0.183 n.s. |
| | History*Selection*Temp | $F_{2,20}$ = 1.359 n.s. |
| Productivity | History | $F_{1,4}$ = 0.531 n.s. |
| | Selection | $F_{1,20}$ = 0.012 n.s. |
| | Temp | $F_{2,20}$ = 48.832 *** |
| | History*Selection | $F_{1,20}$ = 0.096 n.s. |
| | History*Temp | $F_{2,20}$ = 0.286 n.s. |
| | Selection*Temp | $F_{2,20}$ = 0.544 n.s. |
| | History*Selection*Temp | $F_{2,20}$ = 4.374 * |
| Viability | F8 | $F_{1,22}$ = 0.179 n.s. |
| | History | $F_{1,4}$ = 0.100 n.s. |
| | Selection | $F_{1,20}$ = 0.016 n.s. |
| | Temp | $F_{2,21}$ = 39.468 *** |
| | History*Selection | $F_{1,19}$ = 0.437 n.s. |
| | History*Temp | $F_{2,19}$ = 0.531 n.s. |
| | Selection*Temp | $F_{2,20}$ = 0.229 n.s. |
| | History*Selection*Temp | $F_{2,20}$ = 2.043 n.s. |

b)

| Trait | Model parameters | $F_{(df1, df2)}$ |
|---|---|---|
| Age of First Reproduction (A1R) | Temp | $F_{2,22}$ = 49.707 *** |
| | 14 vs. 18 | t.ratio = 9.515 *** |
| | 14 vs. 24 | t.ratio = 7.337 *** |
| | 18 vs. 24 | t.ratio = - 2.178 n.s. |
| Fecundity (F6-8) | Temp | $F_{2,20}$ = 31.417 *** |
| | 14 vs. 18 | t.ratio = - 7.664 *** |
| | 14 vs. 24 | t.ratio = - 2.079 n.s. |
| | 18 vs. 24 | t.ratio = 5.585 *** |
| Productivity | Temp | $F_{2,20}$ = 48.832 *** |
| | 14 vs. 18 | t.ratio = - 6.263 *** |
| | 14 vs. 24 | t.ratio = 3.489 ** |
| | 18 vs. 24 | t.ratio = 9.752 *** |
| Viability | Temp | $F_{2,21}$ = 39.468 *** |
| | 14 vs. 18 | t.ratio = - 0.871 n.s. |
| | 14 vs. 24 | t.ratio = 5.655 *** |
| | 18 vs. 24 | t.ratio = 6.245 *** |

**Table S5.** Thermal reaction norms of the F and W populations, after 31 generations of thermal selection: **a)** ANOVA table and **b)** Tukey HSD tests for the significant factor *Temp*. Data refers to average population values. The F statistic shows the degrees of freedom of effect and error. Significance levels for F values: p> 0.05 n.s.; 0.05>p>0.01*; 0.01>p>0.001**; p<0.001 ***.

a)

| Trait | Model parameters | $F_{(df1, df2)}$ |
|---|---|---|
| Age of First Reproduction (A1R) | History | $F_{1,22}$ = 3.864 n.s. |
| | Selection | $F_{1,22}$ = 9.408 ** |
| | Temp | $F_{2,22}$ = 98.644 *** |
| | History*Selection | $F_{1,22}$ = 0.120 n.s. |
| | History*Temp | $F_{2,22}$ = 1.024 n.s. |
| | Selection*Temp | $F_{2,22}$ = 0.939 n.s. |
| | History*Selection*Temp | $F_{2,22}$ = 0.201 n.s. |
| Fecundity | History | $F_{1,4}$ = 0.727 n.s. |
| | Selection | $F_{1,20}$ = 0.557 n.s. |
| | Temp | $F_{2,20}$ = 29.813 *** |
| | History*Selection | $F_{1,20}$ = 0.216 n.s. |
| | History*Temp | $F_{2,20}$ = 1.170 n.s. |
| | Selection*Temp | $F_{2,20}$ = 0.373 n.s. |
| | History*Selection*Temp | $F_{2,20}$ = 0.164 n.s. |
| Productivity | History | $F_{1,4}$ = 1.560 n.s. |
| | Selection | $F_{1,20}$ = 0.588 n.s. |
| | Temp | $F_{2,20}$ = 35.059 *** |
| | History*Selection | $F_{1,20}$ = 0.128 n.s. |
| | History*Temp | $F_{2,20}$ = 4.079 * |
| | Selection*Temp | $F_{2,20}$ = 2.555 n.s. |
| | History*Selection*Temp | $F_{2,20}$ = 0.507 n.s. |
| Viability | F8 | $F_{1,22}$ = 0.562 n.s. |
| | History | $F_{1,4}$ = 0.947 n.s. |
| | Selection | $F_{1,19}$ = 0.000 n.s. |
| | Temp | $F_{2,20}$ = 32.644 *** |
| | History*Selection | $F_{1,19}$ = 0.055 n.s. |
| | History*Temp | $F_{2,19}$ = 3.078 n.s. |
| | Selection*Temp | $F_{2,19}$ = 1.229 n.s. |
| | History*Selection*Temp | $F_{2,19}$ = 0.424 n.s. |

b)

| Trait | Model parameters | $F_{(df1, df2)}$ |
|---|---|---|
| Age of First Reproduction (A1R) | Temp | $F_{2,22}$ = 98.644 *** |
| | 14 vs. 18 | t.ratio = 12.828 *** |
| | 14 vs. 24 | t.ratio = 11.369 *** |
| | 18 vs. 24 | t.ratio = - 1.459 n.s. |
| Fecundity (F6-8) | Temp | $F_{2,20}$ = 29.813 *** |
| | 14 vs. 18 | t.ratio = - 7.447 *** |
| | 14 vs. 24 | t.ratio = - 1.954 n.s. |
| | 18 vs. 24 | t.ratio = 5.492 *** |
| Productivity | Temp | $F_{2,20}$ = 35.059 *** |
| | 14 vs. 18 | t.ratio = - 5.456 *** |
| | 14 vs. 24 | t.ratio = 2.773 * |
| | 18 vs. 24 | t.ratio = 8.229 *** |
| Viability | Temp | $F_{2,20}$ = 32.644 *** |
| | 14 vs. 18 | t.ratio = - 1.194 n.s. |
| | 14 vs. 24 | t.ratio = 6.100 *** |
| | 18 vs. 24 | t.ratio = 6.619 *** |

**Suppleme**
